# A Cortico-Basal Ganglia Model for choosing an optimal rehabilitation strategy in Hemiparetic Stroke

**DOI:** 10.1101/423863

**Authors:** Rukhmani Narayanamurthy, Samyukta Jayakumar, Sundari Elango, Vignesh Muralidharan, V. Srinivasa Chakravarthy

## Abstract

To facilitate the selection of an optimal therapy for a stroke patient with upper extremity hemiparesis, we propose a cortico-basal ganglia model capable of performing reaching tasks under normal and stroke conditions. The model contains two hemispherical systems, each organized into an outer sensory-motor cortical loop and an inner basal ganglia (BG) loop, controlling their respective hands. In addition to constraint induced movement therapy (CIMT), the model performs both unimanual and bimanual reaching tasks and the simulation results are in congruence with the experiment conducted by Rose et al (2004). Based on our study on the effect of lesion size on arm performance, we hypothesize that the effectiveness of a therapy could greatly depend on this factor. By virtue of the model’s ability to capture the experimental results effectively, we believe that it can serve as a benchmark for the development and testing of various rehabilitation strategies for stroke.

## Introduction

Stroke is considered to be one of the leading causes of disability and mortality worldwide. It could be either ischemic or haemorrhagic in nature. Stroke generally manifests itself as an upper extremity dysfunction with 80 % patients suffering from it acutely and 40% chronically (Cramer, Benson et al. 1997). Sensory and motor deficits resulting from unilateral stroke include difficulty in performing common activities like reaching, grasping and picking up objects. Functionally, hemiparesis is one of the common motor impairments associated with unilateral stroke, after which a more obvious deterioration in the performance of the contralateral arm is observed (Kantak, Jax et al. 2017). Recovery following motor rehabilitation post-stroke primarily depends on the initial severity of paresis (Krakauer 2005) and degree of loss in functionality. It has been reported that the prognosis is good when patients are treated within the first three months following stroke (Wade, Langton-Hewer et al. 1983). Nevertheless, there are reports that show upper extremity recovery ensuing several years after stroke (Carey, Matyas et al. 1993; Yekutiel and Guttman 1993). Hence it has been suggested that progress in functional outcome may be due to neurological repair through cortical reorganization or through compensatory mechanisms (Krakauer 2006).

The ultimate objective of the wide variety of existing rehabilitation protocols like Virtual Reality based rehabilitation (Krichevets, Sirotkina et al. 1995; Jang, You et al. 2005), music therapy (Rodriguez-Fornells, Rojo et al. 2012), Mirror therapy (Oujamaa, Relave et al. 2009), Motor Imagery (Page, Levine et al. 2001; Bajaj, Butler et al. 2015), Motor Imitation (Small, Buccino et al. 2012), Movement observation (Stefan, Cohen et al. 2005), Transcranial magnetic stimulation (rTMS) (Hummel and Cohen 2006), Unilateral muscle strengthening exercises (Patten, Lexell et al. 2004; Daly, Hogan et al. 2005), motor skill learning (Taub, Uswatte et al. 2006), bimanual and unimanual training (Stoykov, Lewis et al. 2009)(e.g., Constraint Induced Movement Therapy (CIMT)), is to enable patients recover from weakness and improve functionality of the arm. However, the problem of determining the best approach for a given stroke patient is still unresolved due to inherent contradictions in some of existing rehabilitation approaches (Hatem, Saussez et al. 2016). For instance, CIMT is primarily based on the repetitive use of the affected arm by restraining the healthy arm (Taub, Uswatte et al. 2006) while bimanual training pairs the healthy arm with the paretic arm to increase the chances of recovery (Stinear, Byblow et al. 2014).

In this paper, we present a computational model of motor stroke affecting upper extremities. We focus on bimanual movements since real world requirements demand coordinated use of both the arms. Recent evidence suggests that mechanisms underlying improvement from bimanual training include the recruitment of ipsilateral corticospinal pathways, increased control from the contralesional hemisphere and normalization of inhibitory mechanisms (Kwakkel, Kollen et al. 2004). An interesting experimental study conducted by Dorian and Rose (2004) has shown that bimanual reaching tasks were more beneficial in reviving the activity of the affected limb post-stroke (Rose and Winstein 2004). A key result of this study is that both aiming distance and bimanual context contributed to the enhanced performance of the paretic arm.

Despite the existence of myriad rehabilitation strategies, the problem of choosing the optimal strategy for a given patient is still unsolved. In this paper we use computational modelling to study two common rehabilitation strategies – bimanual training and CIMT. Whereas bimanual training recommends use of both hands since the normal hands aids in the rehabilitation of the paretic hand, CIMT posits that the normal hand must be arrested to permit optimal rehabilitation of the paretic hand. The two strategies are obviously mutually contradictory. Which strategy is more suitable in a given situation? We address this question using an elaborate computational model of the cortico-basal ganglia network.

Our model is designed to perform both unimanual and bimanual reaching under different task conditions. The model consists of an outer cortical loop and an inner basal ganglia loop. Performance is evaluated in terms of peak resultant velocity (PRV) and reaching error. The entire system drives a two-link arm model engaged in targeted reaching movements. The basal ganglia loop primarily drives motor learning with the control gradually passed on from the basal ganglia to the motor cortex as learning progresses. Two copies of the entire system, with appropriate coupling, are used to simulate bimanual reaching. The model is able to explain reaching behaviour under normal and hemiparetic stroke conditions and is also able to capture the bimanual reaching experiments of Dorien and Rose (2004). The model is also operated under CIMT conditions. A comparison of the effect of the two apparently contradictory strategies on model performance pointed to an interesting resolution: bimanual reaching is found to be more beneficial for smaller lesion sizes whereas CIMT is more effective for bigger lesion sizes.

The outline of the paper is as follows. Section 4 describes the model equations and the task setup. Section 2 describes the model results related to unimanual and bimanual reaching. A comparison of recovery from simulated stroke following CIMT and bimanual reaching was also presented. Section 3 presents a discussion of the results and scope for future work.

## Results

### 1. Training the outer motor cortical loop under normal conditions

The training schema of the entire cortico-basal ganglia model is shown in figure 1.

**Fig.1:**
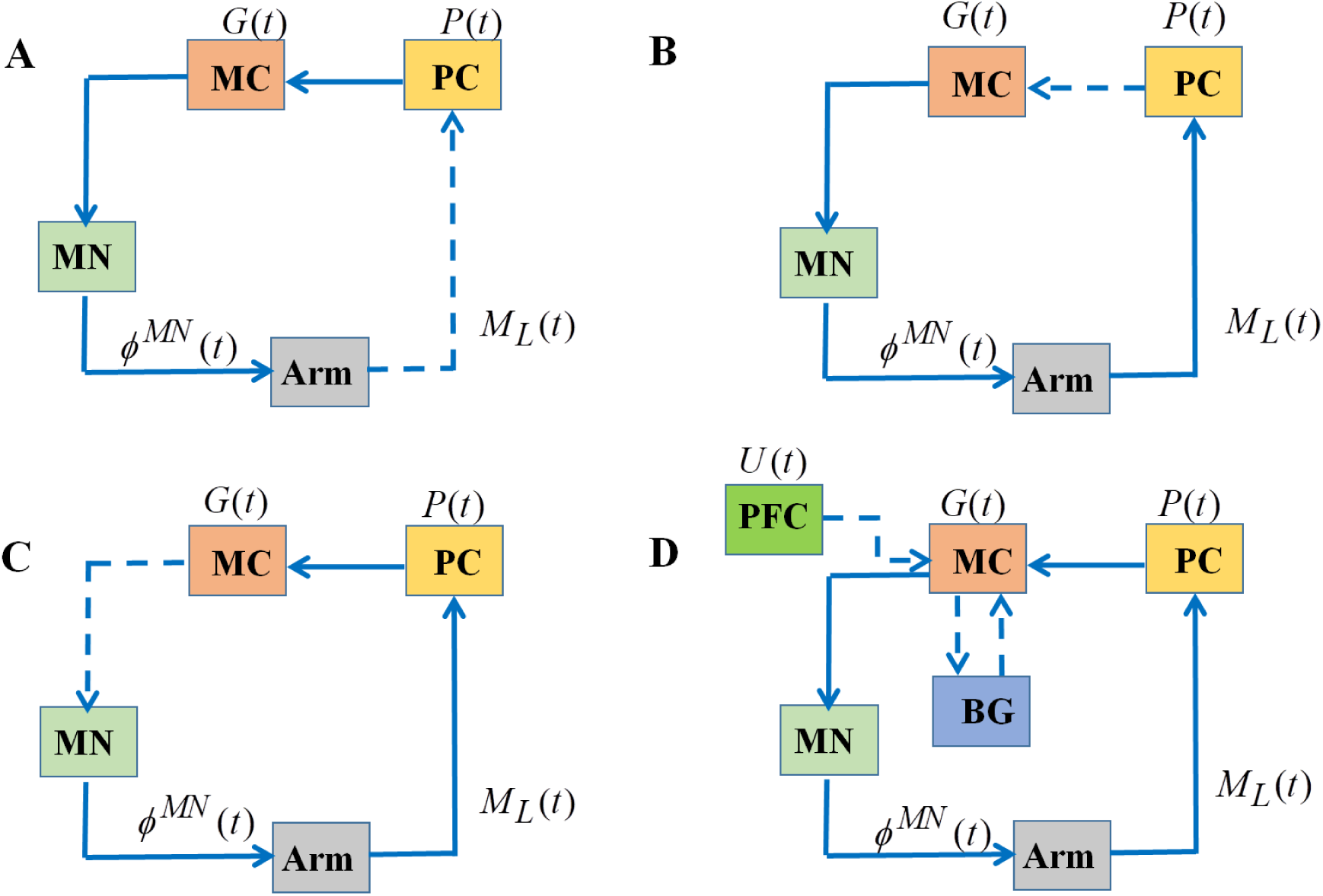
The training schema in the cortico-basal ganglia model. (**A**) training the Arm to PC connections (**B**) training the PC to MC connections, closing the loop by (**C**) training the MC to MN weights. Then the BG module is introduced and the PFC to MC connections are trained (**D**). In every figure, the dashed arrows indicate the connections that are being trained.

#### Training the weight connections between PC and MC

- Random activations of the agonist-antagonist muscle pairs will place the arm in ‘n’ different configurations with each configuration corresponding to a muscle length vector (M_L_). This will then serve as feature vectors to train the SOM of the proprioceptive cortex using the standard SOM algorithm (Muralidharan, Mandali et al. 2018).
- The SOM response of the PC layer is then projected to MC SOM. Since every node in PC is connected to every node in the MC, this accounts for low dimensional representation of the sensory input to the motor cortex.

#### Training the weight connections between MC and MN

- To begin with, a random activation vector is presented to the arm which subsequently activates it, thereby setting the arm in an equilibrium configuration.
- In an ideal state, the flow of this sensory information via PC to MC and then back to MN, must give rise to muscle activation equal to the random activation vector that was presented to the arm at the outset. However, since no system is ideal, the connection between MC and MN is trained by pushing the actual MN activation towards the desired activation, using equation (**22**).

### 2. Mapping the arm configurations

The sensory motor loop is tested at the level of motor cortex to determine the range of arm movements in the workspace. A direct Gaussian current is given to the MC of size 25×25 based on the following equation:

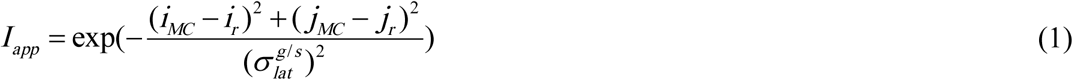

where, *I_app_* is the input Gaussian current, i_MC_ and j_MC_ represent the MC nodes and i_r_ and j_r_ is the random node at which the current is centred.

It is observed from figure 2(B) that, most of the positions in the 2D workspace are accessible by the arm. Also, on analogizing, we found that the activity generated probing the MC and the subsequent activity generated by signal flow via PC to MC is the same.

**Fig 2.**
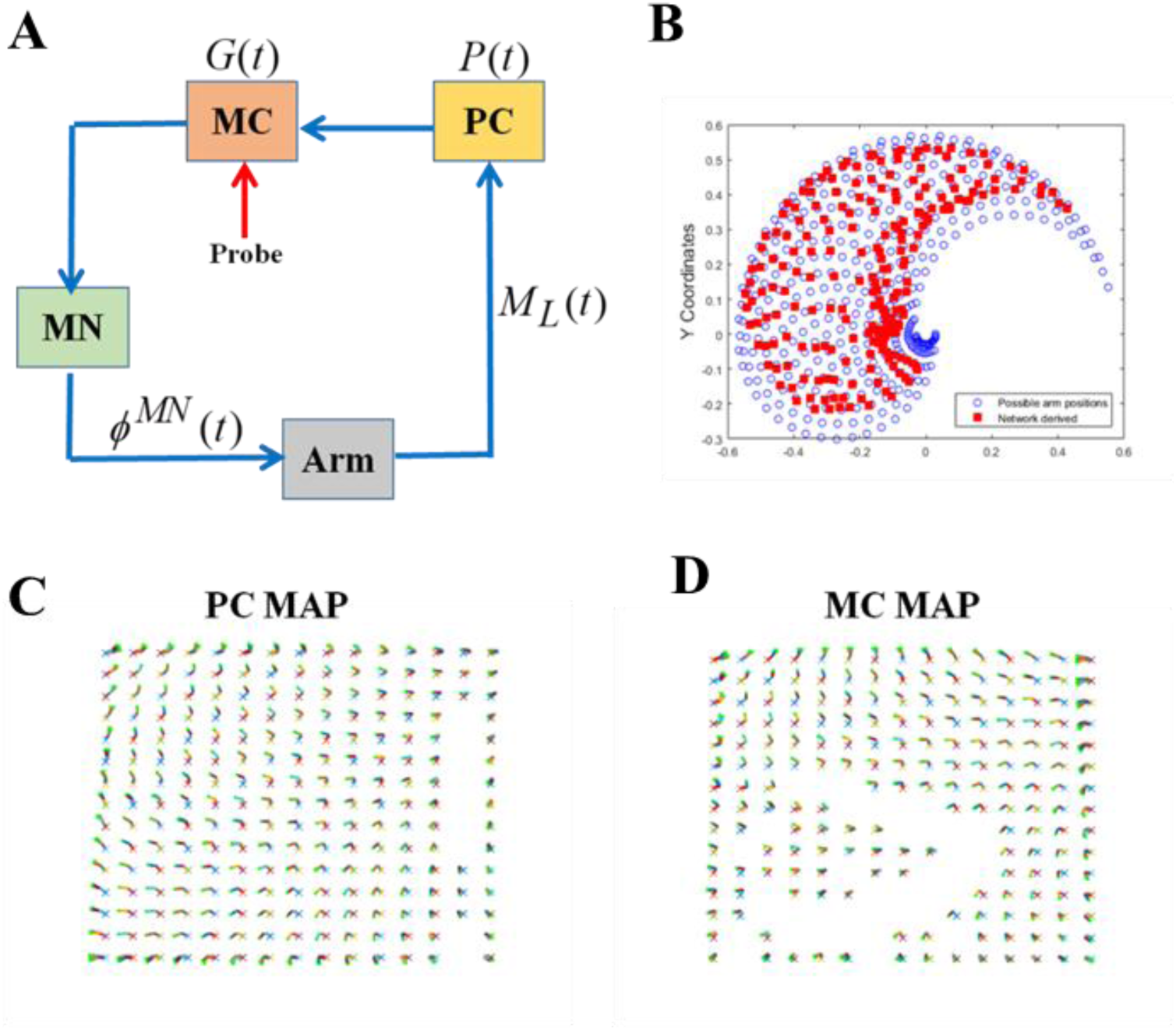
Sensory and Motor Maps. (**A**) The sensory-motor loop is probed at the level of the MC. (**B**) Mapping of the end effector positions approximated by the network is compared to all possible positions in the arm workspace. (**C,D**) The joint configuration maps formed for both the PC and the MC layers respectively.

### 3. Reaching movements of the arm

During early stages of learning, movements of the arm are predominantly directed by the BG output which in later stages is combined with the increasing contribution of the PFC (Muralidharan, Mandali et al. 2018). The PFC activity that codes for the target location defines the activity that the motor cortex must evolve to in order to reach the target. The complex oscillations of the STN-GPe system enable adequate exploration of the arm in the workspace. Every time the arm moves within a distance of ‘r’ units from the target, it is considered to have accomplished its reaching task and in turn leads to training of the weight connections between PFC and MC. It is to be noted from the figure 3(A,B) that, when the PFC input to the MC and the resultant MC activity are identical, it refers to the fact that the network has learnt to approximate the activity needed to reach the target. Currently in the simulation, there are a total of 50 trials, out of which the first 30 correspond to training period of the arm and the last 20 correspond to testing the performance of the arm. Each trial is initialized from a starting position and the goal location is kept constant throughout all trials. For every successful reach, the magnitude of the PFC contribution is increased, while training of the value function and PFC to MC connections occur in parallel. Since our model is bimanual, a trial is terminated only when both arms are successful in reaching their respective targets.

**Fig 3.**
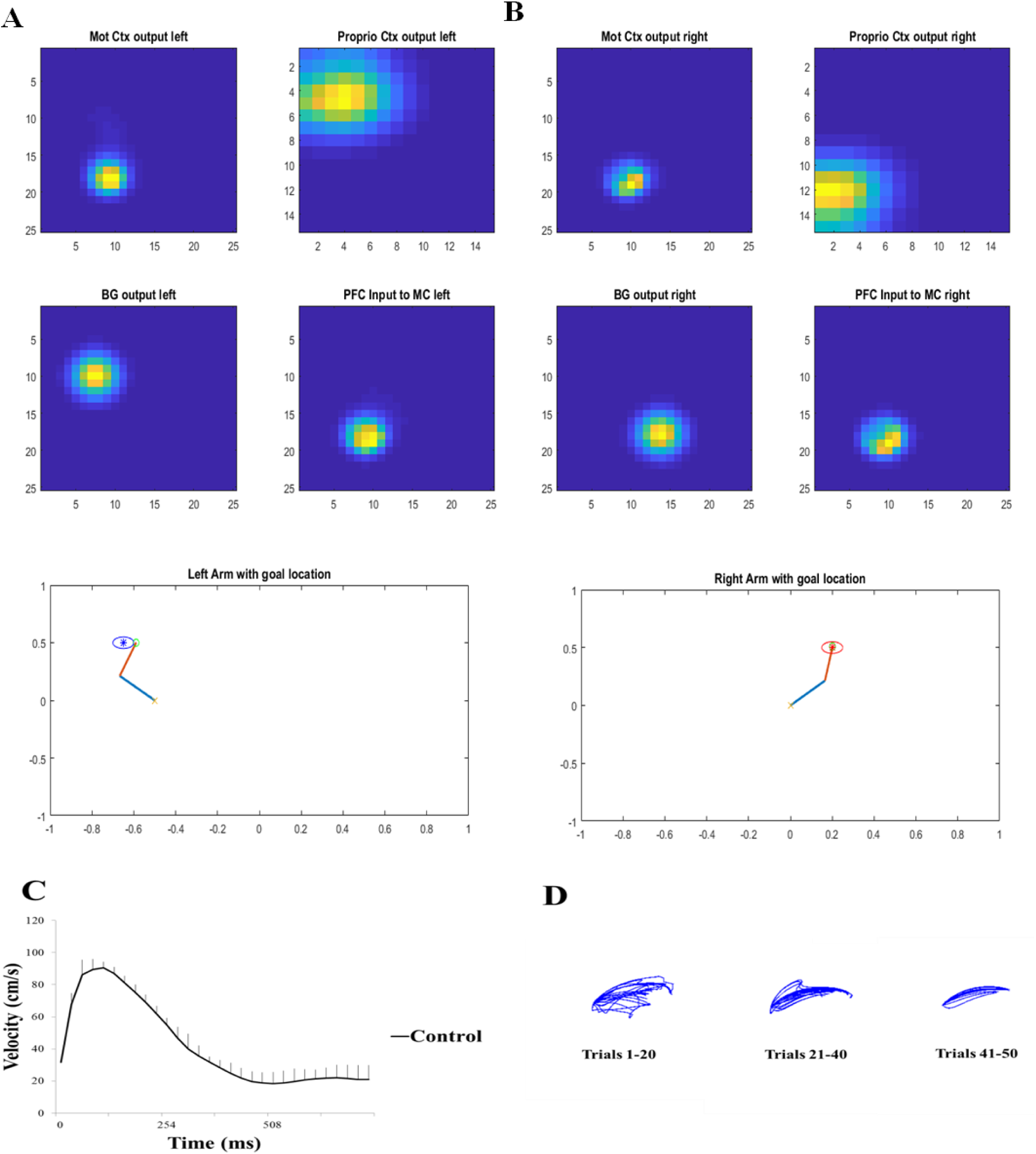
Reaching behaviour in healthy controls. (**A, B**) The network output of the left and right arm while performing the reaching task independently and the activities of multiples areas in the model. (**C**) The velocity profile of the right arm during a reach; this will serve as control to compare before and after lesion introduction (**D**) The end effector trajectories obtained in the case of the control for reaching a single target across trials as the learning of the PFC to MC connections (W_PFC→MC_) takes place.

**Fig 4.**
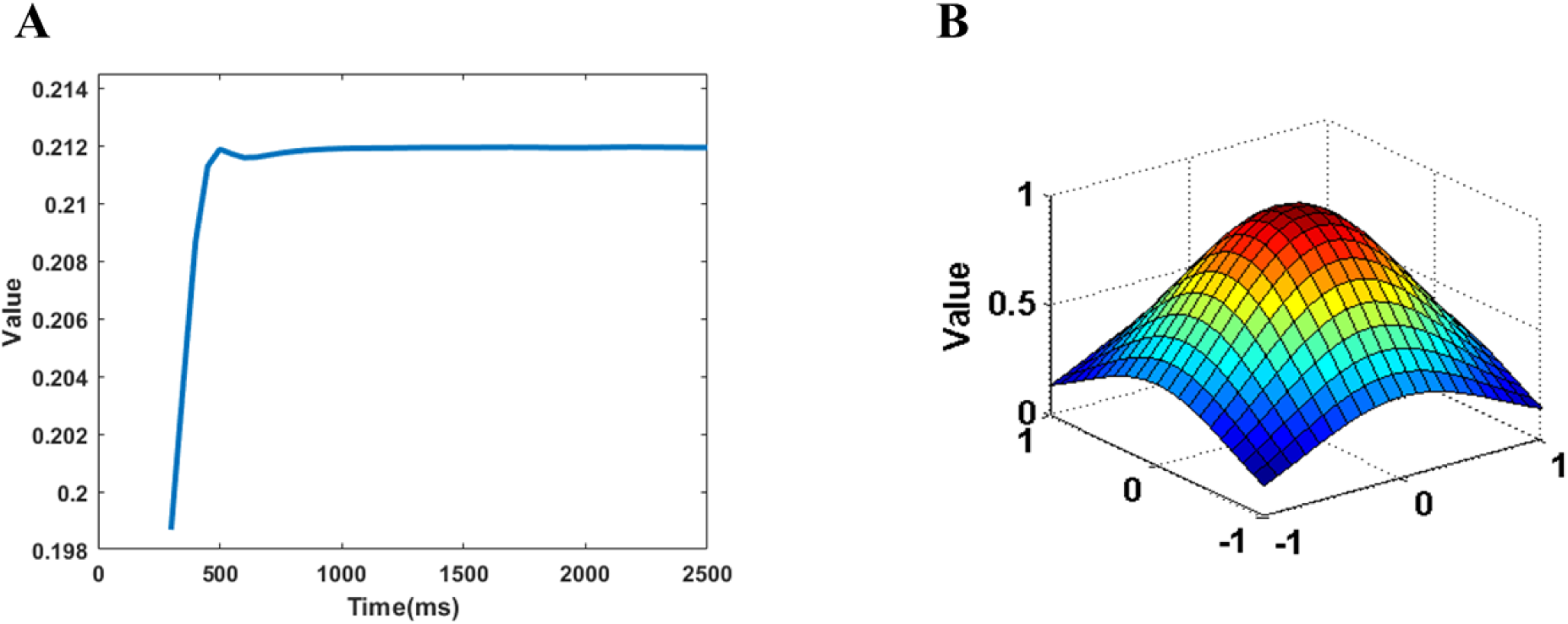
Value function. **(A)** Value building over time. **(B)** The value function is maximum at the target location; in this case the target is at (0,0).

From figure 3(D), it is observed that the end effector trajectories of the arm movements, become less erratic and more smooth, indicating the decrease in hand path variability as learning progresses. Furthermore, on analysing the velocity profile of the arm performance in control conditions, we found that it exhibits a bell-shaped profile (Muralidharan, Mandali et al. 2018).

### 4. Training the Value function in the Basal Ganglia

The Basal Ganglia (BG) network in the model consists of the following components – a striatum containing D1-R and D2-R expressing medium spiny neurons, a globus pallidus external (GPe), a globus pallidus internal (GPi), a subthalamic nucleus (STN) and a thalamus. Consistent with our previous models, the BG network is trained using reinforcement learning algorithm with dopamine coding for a temporal difference error signal. The value computed at every time instant t (eqn **23**), is given as the input to the BG.

The value difference signal (eqn **24**) is then carried via nigrostriatal connections to the striatum where they modulate the activity of D1-R and D2-R expressing neurons according to eqns. (**25, 26**).

In eqns. (**25, 26**), *t*_D1_ and *t*_D2_ act as the threshold values of the Direct (DP) and Indirect pathway (IP) respectively. The DP acting via the striatum, GPi and thalamus, is responsible for movement activation, while the IP acting via the striatum, GPe, STN and the thalamus, is responsible for movement inhibition. The selection between the two pathways depends on the input to the striatum.

The value function implemented by a multilayer perceptron comprises of an input layer, a hidden layer and an output layer. PC activity at times (t) and (t-1) combined with the PFC activity serves as the input to the network and is utilized to compute the TD error. Training of the network is performed by means of error backpropagation.

The value function computed by the BG has its maximum value at the target/goal. Thus, the BG performs a stochastic hill climbing over the value function and in doing so, enables the arm to reach the goal/target. During the initial periods, the movements of the arm are governed by value gradient information; hence its behaviour tends to be more exploratory in nature. As the trials progress, the BG becomes more adept at finding the maximum of the value function thereby driving the arm to make more direct movements, whose activity is now dominated by the PFC.

### 5. Simulating intra-cortical connectivity

The bimanual aspect of the model has its seed at the level of the respective motor cortices of the two arms. The MCs communicate by means of a “coupling factor (*ε*)” where, the product of *ε* and MC activity 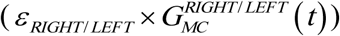 of the ipsilateral hemisphere is added with the input current to the MC CANN 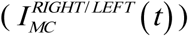 of the contralateral hemisphere. Based on eqn. (**18**), this coupling is mathematically represented by,

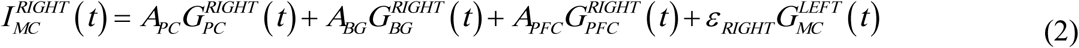

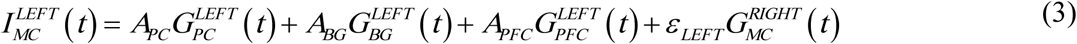

Based on the coupling strengths, the reaching task conditions could be categorized into:

a. **Unimanual condition:** Here the values of both *ε_RIGHT_* and *ε_LEFT_* are equated to zero thereby signifying that the arms function in an independent manner.
b. **Bimanual condition:** Here the coupling values of *ε_RIGHT_* and *ε_LEFT_* are non-zero and are optimized to fit the data.
c. **Constraint induced movement:** Here the coupling values have the same range as in bimanual condition. However in this case, unlike bimanual, 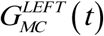 will remain constant denoting the dormant nature of the constrained, unaffected (left) arm.

The velocity profiles (Fig. 3C) of the aforementioned task conditions when performed in the absence of the lesion, serve as control data for comparison with performance of the model post-stroke.

### 6. Simulating Hemiparetic Stroke in the model

To simulate and study hemiparetic stroke, we incorporated “lesion” of size n x n in the right MC of our model by suppressing the activity of a fixed number of nodes in g_MC_ of the CANN as follows:

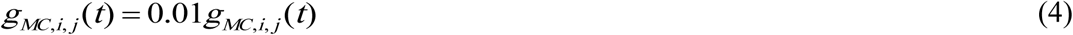

where,

g_MC_ is the state of neurons in MC CANN at time ‘t’.

(i,j) corresponds to a 2D array in g_MC_ that is used to define the lesion in MC.

a and b define the location of the lesion in the MC.

### 7. Performance of the arm post-stroke simulation

To test changes in the kinematics of the arm after introducing lesion, we estimated the velocity profile of the affected arm while it was performing a reaching task. The network was trained to reach a single target followed by introduction of lesion in the corresponding MC. It can be noticed from fig. 6C that there is a significant decrease in the velocity of the affected arm when compared to its normal counterpart.

### 8. Training the outer motor cortical loop under lesion conditions

#### Retraining PC to MC connection

Once a lesion is introduced, PC to MC connection is retrained such that the winner node corresponding to the end effector position of the arm is picked from the neighbouring nodes that lie outside the lesion area. The winner node is the one whose weight is the least distant from the given input vector. Also, instead of random initialization of SOM weights, weights trained under normal conditions are used to retrain the connections between PC and MC.

#### Retraining MC to MN Weights

MC to MN weights are retrained in order to maintain consistency with respect to retraining the cortical loop once a lesion is introduced. Therefore, the weight updation is given as,

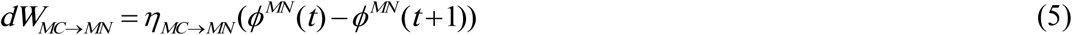

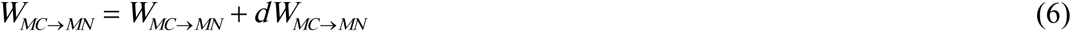

Where,

*η*_*MC*→*MN*_ is the learning rate

*dW*_*MC*→*MN*_ is the weight update

*ϕ^MN^* (*t*) and *ϕ^MN^* (*t* +1) are the MN outputs computed from motor cortical activities collected at t^th^ and t+1^th^ time steps when the arm is closest to its goal. The recovery of the lost end effector positions in the workspace after retraining the cortical loop is as shown in fig. 5C.

**Fig 5.**
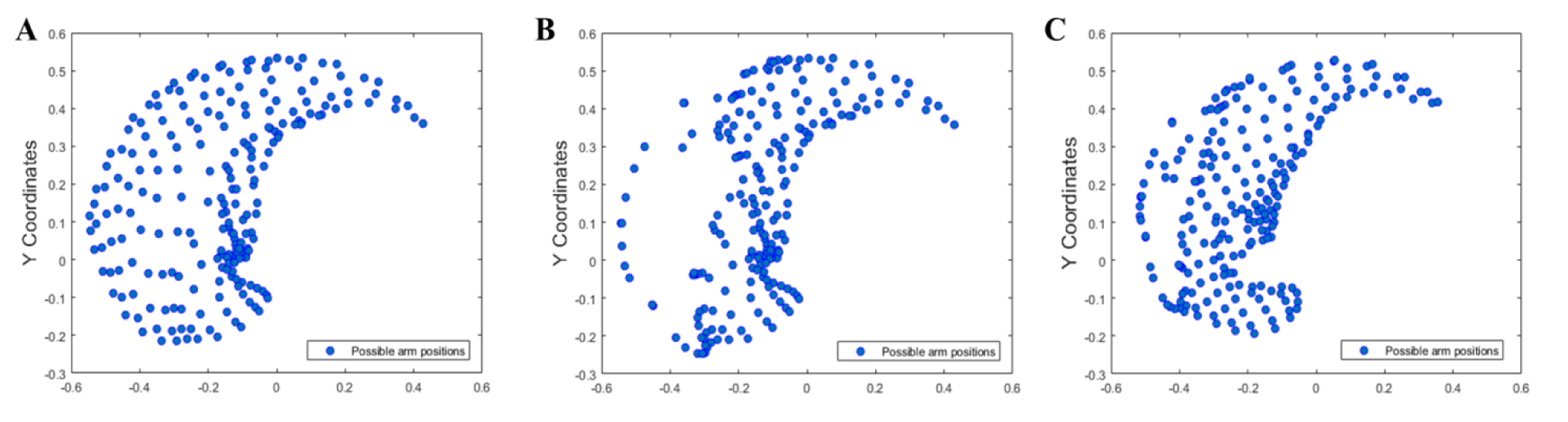
Mapping of the end effector positions approximated by the network. All possible end effector positions of the arm in the given workspace are mapped under three different conditions **(A)** normal(control) condition **(B)** stroke condition and **(C)** retraining the cortical loop post stroke

**Fig 6.**
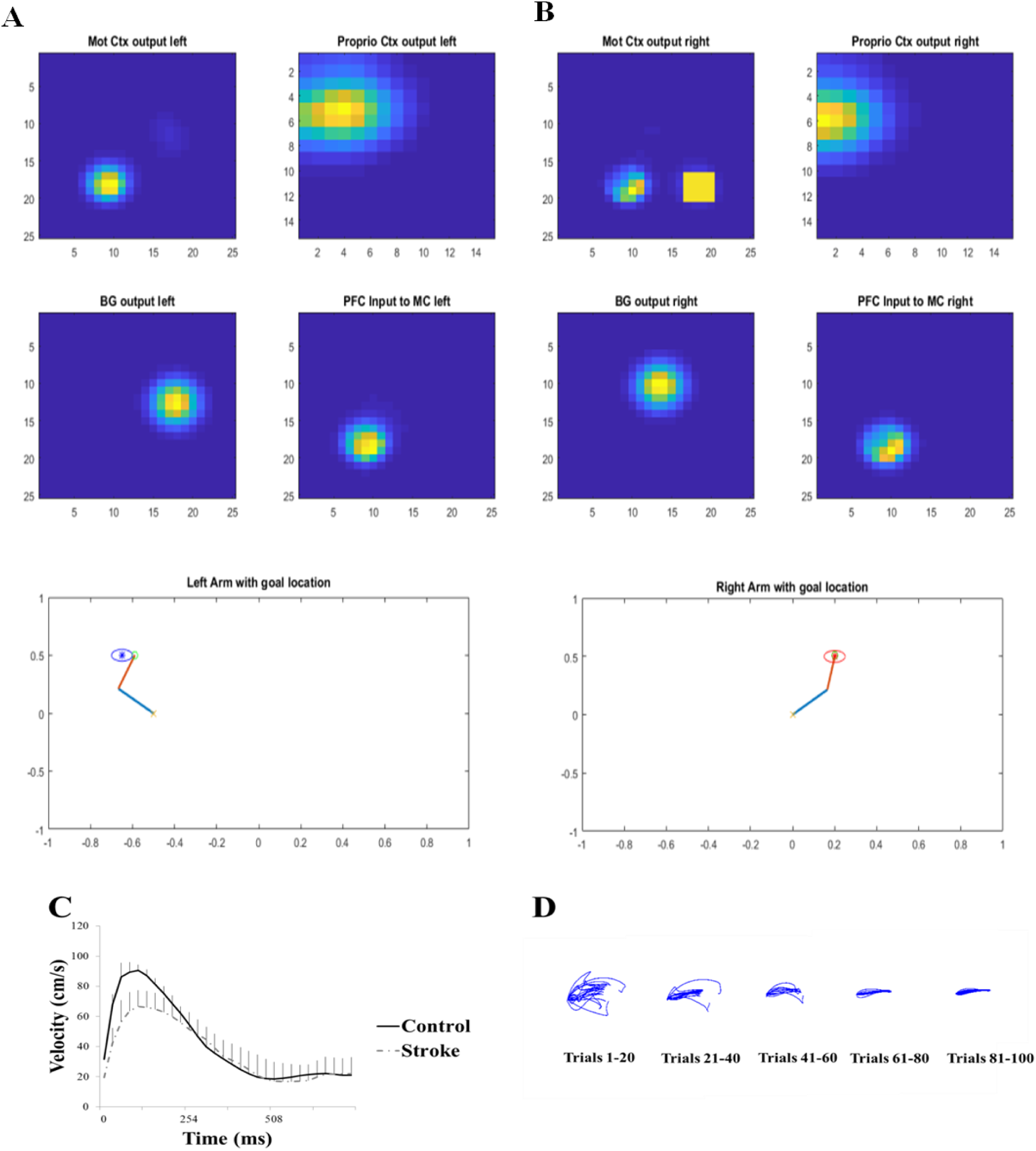
Reaching behaviour in stroke. (**A, B**) The network output of the left and right arm while performing the reaching task independently and the activities of multiples areas in the model. (**C**) The velocity profile of the right arm (now paretic) during a reach. The yellow square in MC right denotes the presence of lesion. (**D**) The end effector trajectories of the paretic arm obtained for reaching a single target across trials as the learning of the PFC to MC connections (W_PFC→MC_) takes place.

### 9. Experimental study – Rose and Winstein (2004)

Rose and Winstein (2004) investigated the feasibility of bimanual training protocol for post-stroke rehabilitation. They tested the performance of the upper extremities under unimanual and bimanual conditions in three different reaching paradigms. A total of 30 stroke patients, alongside 30 healthy people who served as controls, were involved in the study. The participants were asked to reach forward rapidly and aim with one hand (unimanual) or both hands (bimanual) and hit switches mounted on LED targets in response to an LED signal.

Their initial study examined a spatially symmetric forward aiming movement of the arms. They found that, in unimanual condition, the non-paretic arm exhibited a higher Peak Resultant Velocity (PRV), than in bimanual condition, where it was paired with the paretic arm. On the contrary, the paretic arm exhibited a higher PRV when it functioned bimanually as opposed to unimanual condition. Interestingly, this discrepancy was observed only in stroke patients and not in controls.

Rose and Winstein (Rose and Winstein 2004) further extended their study by inducing spatial disparity in reaching distances which would require proper co-ordination among both the arms. Similar to their previous case, the participants had to aim and hit switches in response to an LED signal, only that the arms moved to separate targets. This experiment was classified into two based on the level of difficulty of the task assigned to paretic arm. The first type was called “congruent aiming”, where the paretic arm moved to a “near” target and the non-paretic arm moved to a “far” target while the second type known as “incongruent aiming” followed the converse of the congruent task setup (fig. 7).

**Fig 7.**
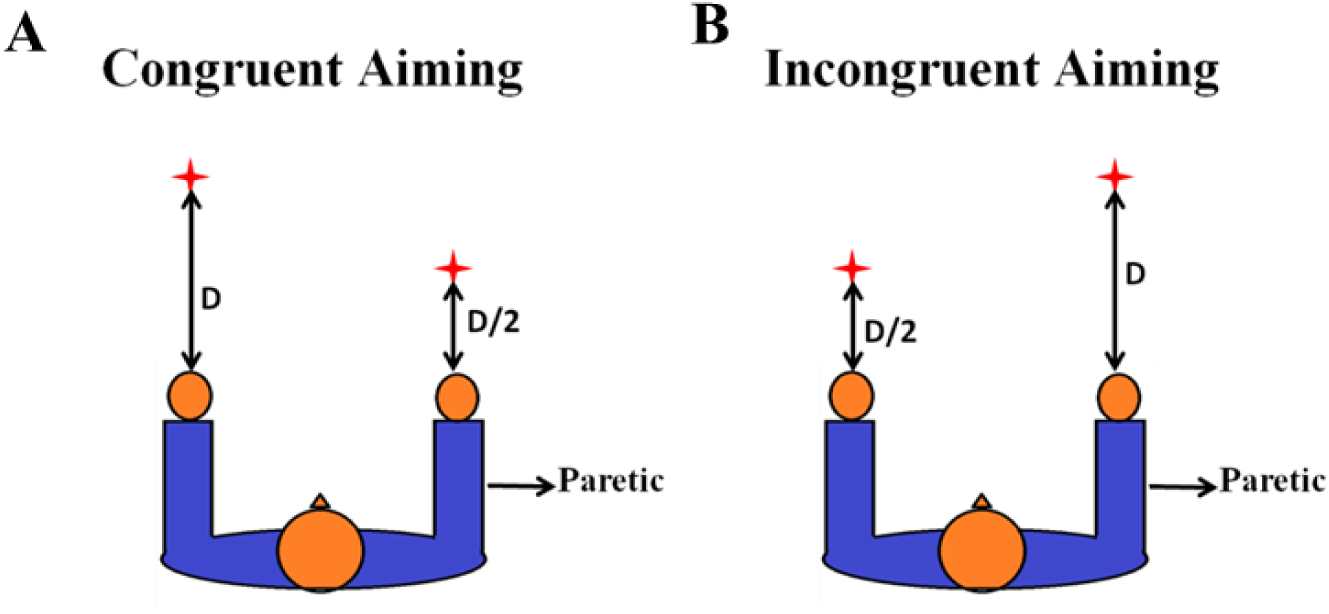
Asymmetric target aiming. (**A,B**) Congruent and Incongruent aiming setups in Rose and Winstein’s experiment. ‘D’ refers to the aiming distance. In congruent condition, target for the paretic arm was located at half the distance of target location of the normal arm. In incongruent aiming, the paretic arm had the farther target to reach.

The “near” target is placed at 50% of the distance of the far target as shown in fig. 7. It was observed that the non-paretic limb exhibited a prolonged movement execution time accompanied by a decrease in its PRV in bimanual condition, than that in unimanual condition. Also, there was an increase in the paretic limb PRV in bimanual condition when compared to unimanual condition. However, the enhanced paretic limb PRV was seen only with respect to incongruent aiming which further led them to compare paretic limb PRV in unimanual conditions for the near and far aiming tasks to verify if aiming distance alone facilitated such an increase (Rose and Winstein 2004).

The paretic limb PRV was similar in both unimanual aiming conditions thereby suggesting that both paretic aiming distance and constraints of bimanual coordination are prerequisites to improve performance of the paretic arm.

### 10. Model performance on the reaching tasks

#### Symmetric Aiming

We simulated the symmetric aiming task in the model by providing each arm with its respective target, placing them at spatially symmetric locations as shown in fig. 8D. In the first case, the arms are allowed to perform the reaching under unimanual aiming conditions where ε=0. This is followed by testing the arm under bimanual conditions using both inhibitory and excitatory coupling (ε < 0 and ε > 0). For each arm, its velocity at every instant in time is recorded and averaged to find the maxima of the velocity profile which is referred to as the PRV.

**Fig.8:**
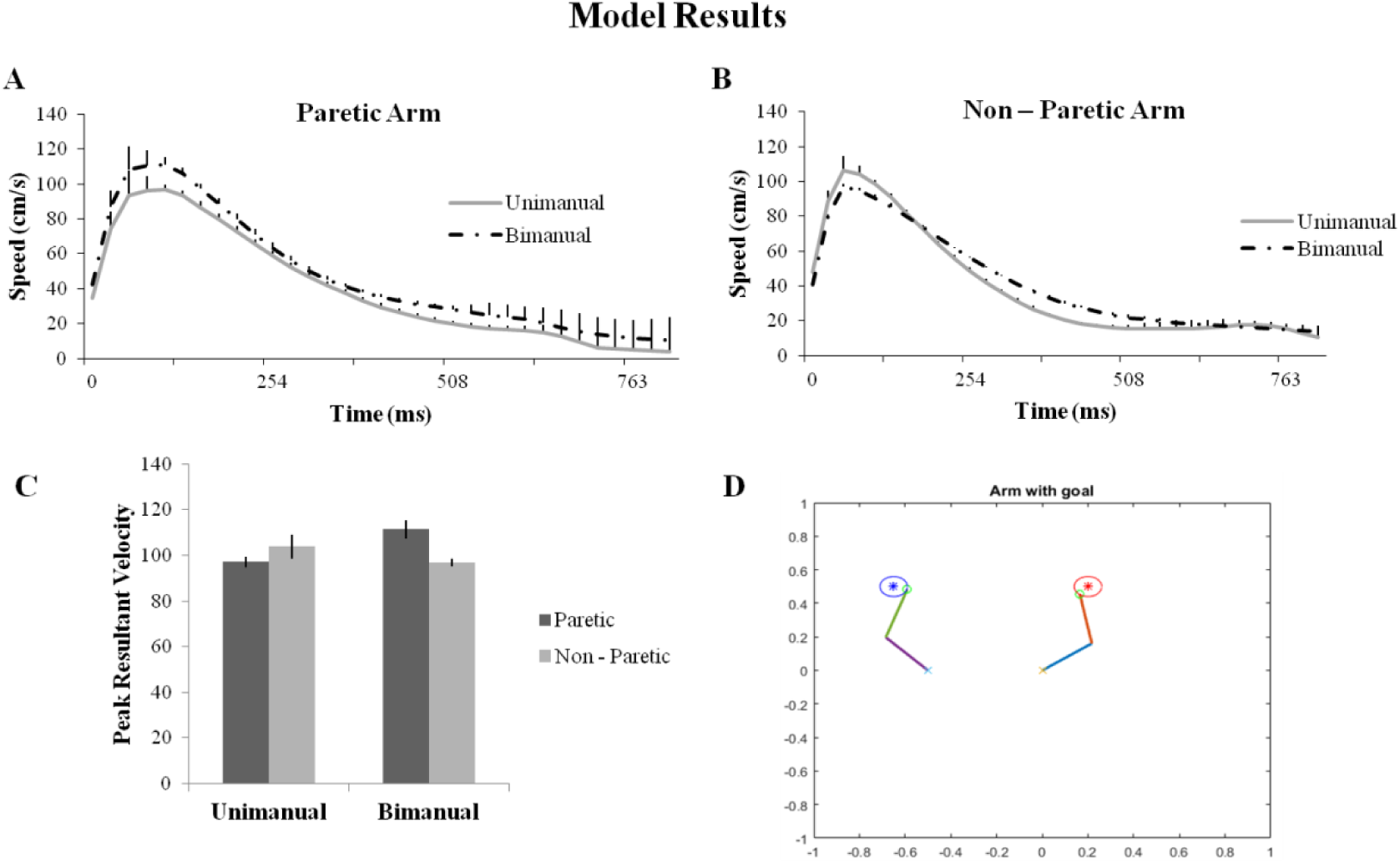
Model performance in symmetric aiming. (**A, B**) Velocity profiles of the paretic and the non-paretic arm under unimanual and bimanual conditions obtained from the Model. (**C**) The bar graph of the Peak resultant velocities (from model). (**D**) Snapshot of the simulation of the arms reaching the targets in symmetric aiming task.

It is observed that under unimanual conditions, the non-paretic arm (left arm in the model) shows a greater PRV than the paretic arm (right arm in the model). Similarly, it is found that during bimanual training of the arms to accomplish the target, the paretic arm showed a significant increase in its PRV whereas the non-paretic arm showed a decrease in its PRV when compared to unimanual training as shown in fig. 8A and B. This was achieved by using the coupling factor where there is excitatory influence from the paretic to the non-paretic arm and inhibitory influence from non-paretic to the paretic arm.

#### Congruent and Incongruent Aiming

The second experiment conducted by Rose and Winstein required movement of the upper extremities to “near” and “far” targets. Based on this spatial disparity, the tasks were divided into congruent and incongruent aiming tasks. In congruent aiming, the paretic arm had a near target and hence an easy task to accomplish, whereas the non-paretic arm had a far target to reach. The near target is placed at 50% of the distance of the far target as shown in fig. 9B. We simulated this task setup in the exact same manner and allowed the model to perform the task. The PRV’s of both the arms are recorded under unimanual and bimanual conditions. It is found that under unimanual conditions, the non-paretic arm had a higher PRV than the paretic arm with, no significant change in the PRV of the paretic arm in the bimanual case. However a decrease in the velocity of the non-paretic arm is observed, when it is paired with the paretic arm (i.e. bimanual) (Fig. 9A).

**Fig.9:**
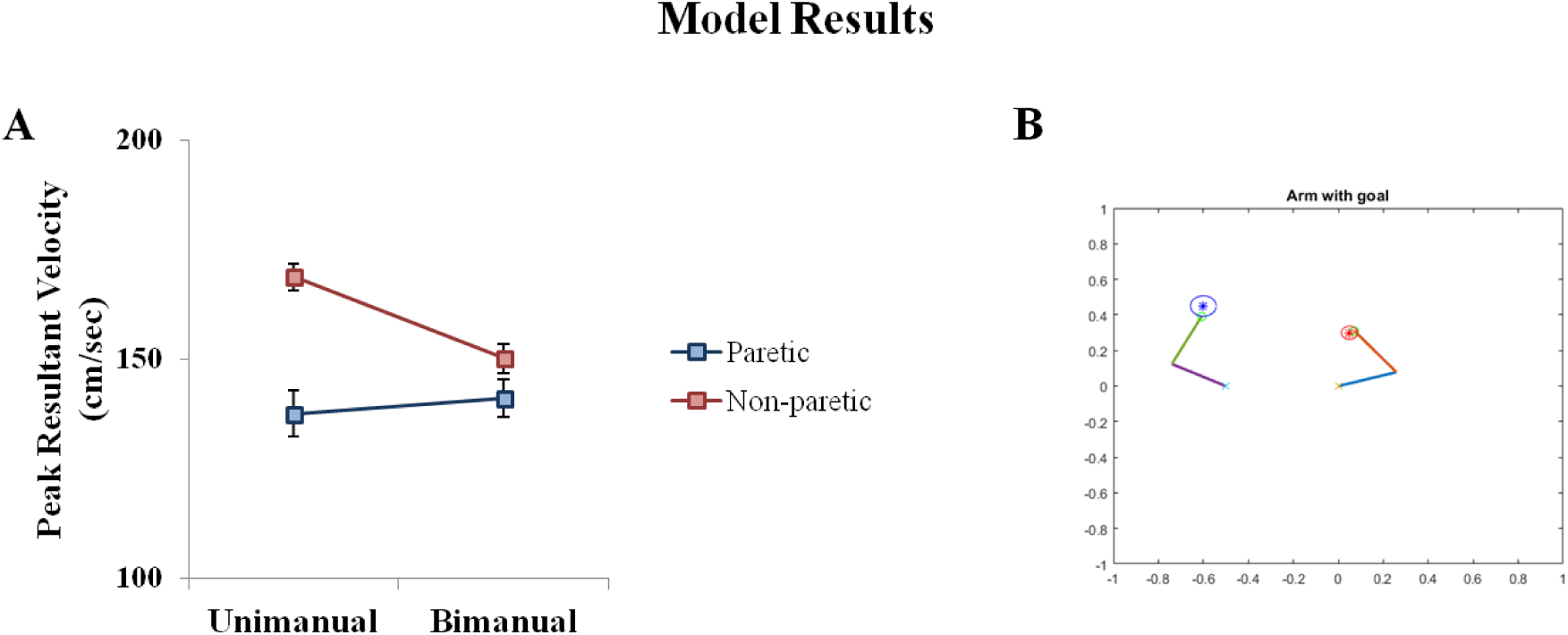
Model performance in congruent aiming. (**A**) Peak resultant velocities of the paretic and the non-paretic arm under unimanual and bimanual conditions obtained from model (**B**) Snapshot of the simulation of the arms reaching the targets in congruent aiming task.

The incongruent aiming task observes the converse experimental setup as that of the congruent aiming task. Here, the paretic arm (right) has a far target (difficult task) to accomplish whereas the non-paretic arm (left) has a near target to accomplish as shown in fig. 10B. A similar recording of the PRV is performed as mentioned in the earlier studies. It is found that the PRV of the non-paretic arm is higher than the paretic arm under unimanual conditions. But in the bimanual conditions, the paretic arm shows a significant increase in its velocity whereas the velocity of the non-paretic arm decreased greatly as shown in the graph (fig. 10A). The results obtained in all the three aforementioned task setups are identical to those obtained by Rose and Winstein (Rose and Winstein 2004).

**Fig.10:**
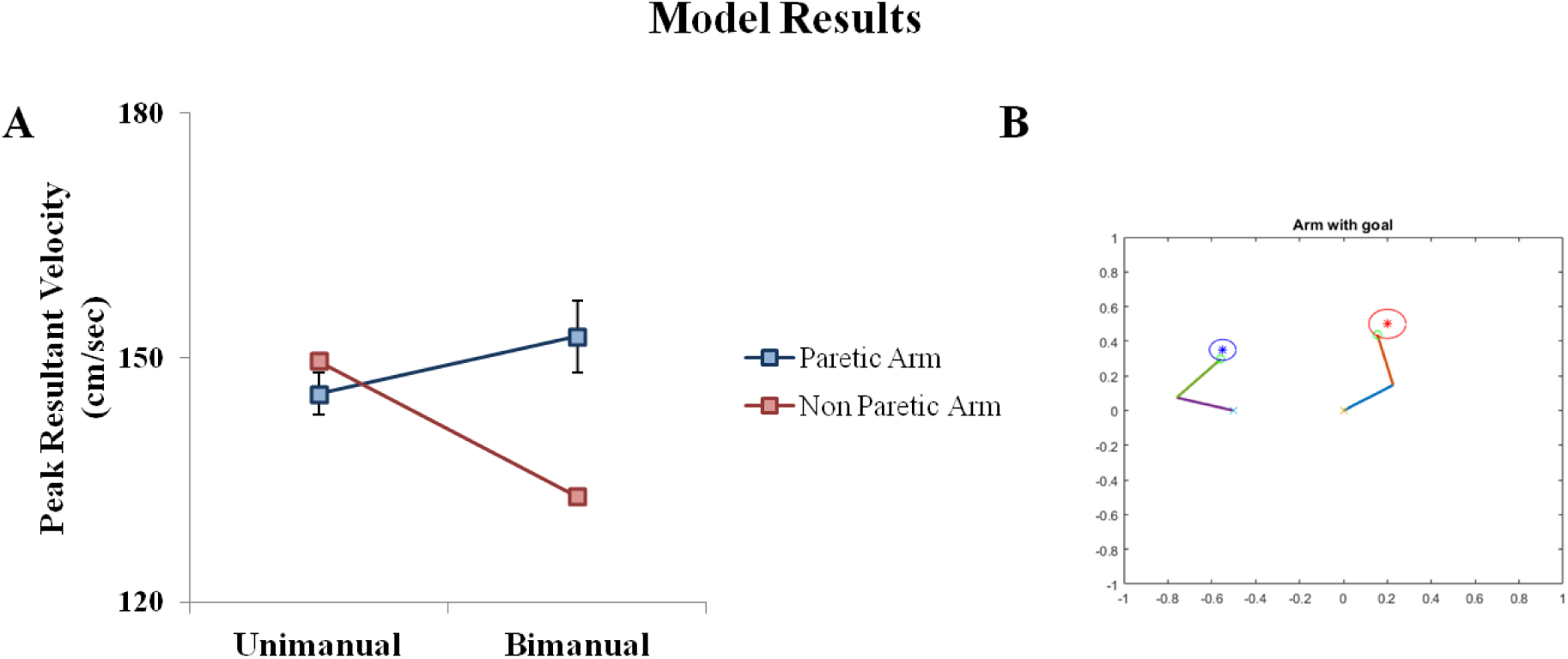
Model performance in incongruent aiming. (**A**) Peak resultant velocities of the paretic and the non-paretic arm under unimanual and bimanual conditions obtained from model (**B**) Snapshot of the simulation of the arms reaching the targets in incongruent aiming task.

### 11. Aiming conditions Vs lesion size

In this part, we study the effects of different lesion sizes of the MC on reaching behaviour. One of the several factors inducing a change in reaching performance could be the extent of damage in the corresponding hemisphere after stroke. The model provides a framework to test this effect and also serves as a tool to compare different interventions currently available to find the best possible one for patients in order to help them regain control. It becomes important to provide a forum that can reconcile different strategies since most of them are inconsistent with each other in relation to their techniques.

Here, we record the reaching error of the arm i.e. the minimum distance of the arm from its target location over lesion sizes starting from 1×1 to 7×7 for CIMT, unimanual and bimanual reaching tasks. The eye of the lesion is exactly on the neuron that gets activated whenever the arm reached its target.

In addition, we tested the model performance in two stages of stroke namely, chronic and acute. In chronic stroke, the MC was trained with lesion and then tested. This could be equated to a condition where therapy was provided a few months after the incidence of stroke. While in the acute stroke condition, the MC was trained normally and tested right after introducing the lesion without any further retraining. This could be analogous to a condition where therapy was provided soon after the incidence of stroke.

Upon analysis, for chronic stroke, both unimanual and bimanual conditions of the task completion is observed. It is found that for lesion sizes ranging between 1×1 and 5×5, bimanual training proved to be more beneficial in terms of reduced reaching errors of the paretic arm, whereas lesion studies beyond 5×5 size (6×6 and 7×7) unimanual training of the arm proved to be beneficial.

For CIMT, it is observed that the reaching error decreases as the lesion size increases when compared with bimanual condition. The value of the reaching error obtained for lesion sizes above 5×5, i.e. 6×6 and 7×7 is comparable to that obtained in unimanual condition. Thus, for higher lesion sizes, CIMT also proves to be beneficial when compared with bimanual condition.

For acute stroke, the value of reaching error is very low in bimanual condition up to a lesion size of 6×6. However, for 7×7 the error rises to a significant level (fig. 11). The opposite pattern is observed for unimanual condition and CIMT. For these, the value of reaching error reduces for larger size of the lesion. This is similar to the trend we observed for chronic stroke.

**Fig.11:**
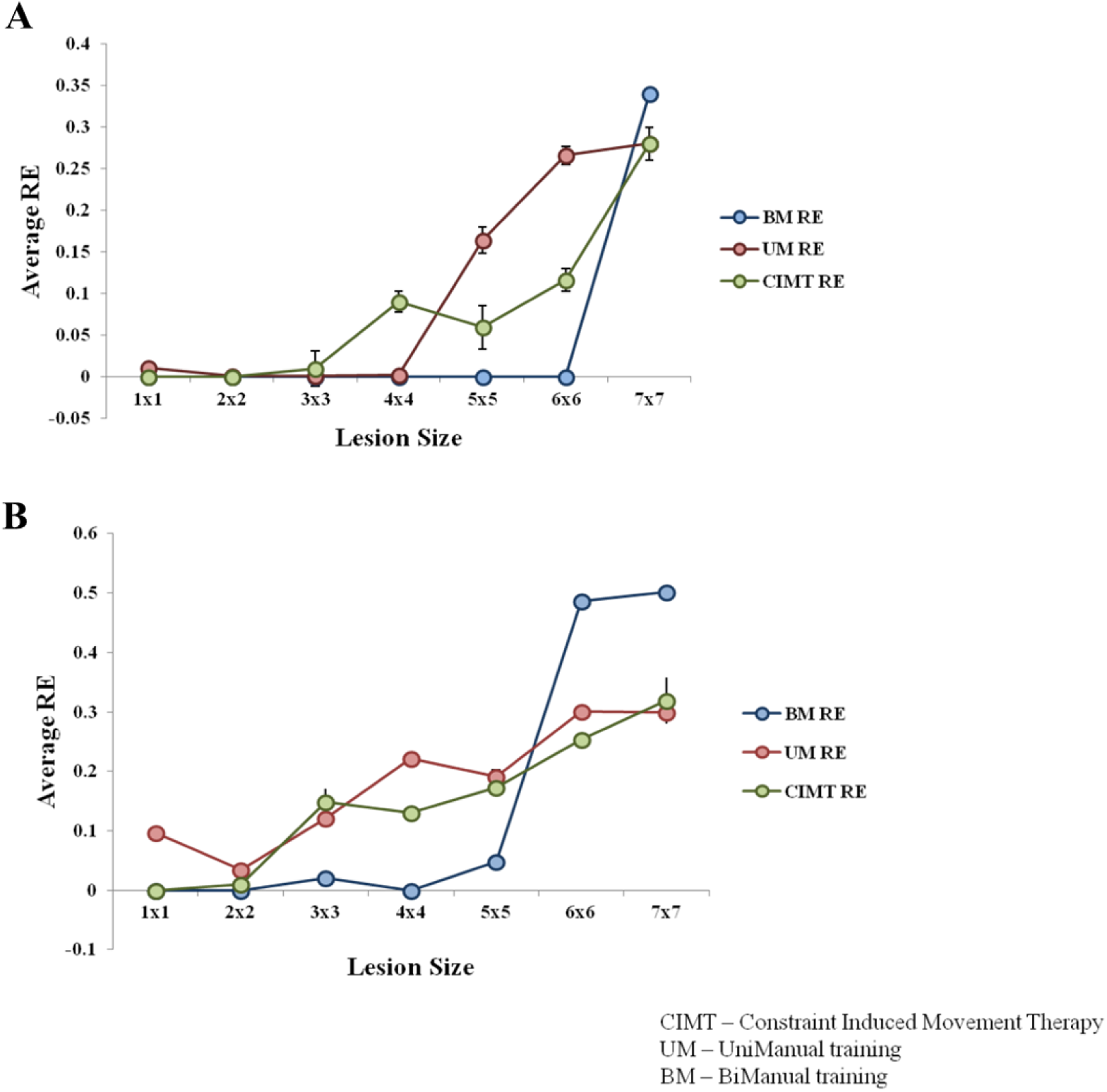
Model performance in varying lesion sizes. (**A, B)** The average reaching error obtained from the model under unimanual, bimanual and constraint induced training conditions plotted with respect to lesion size in acute and chronic stroke respectively.

## Discussion

We propose a biologically plausible model that can perform both unimanual and bimanual reaching in simple task setups. The model is primarily characterized by two loops: the sensory-motor cortical loop and the cortico-basal ganglia loop. This combined system controls targeted reaching in a two-link arm model moving on a two-dimensional plane. Two copies of the entire system are used to simulate bimanual reaching. We consider only motor stroke affecting the upper extremities in this model and stroke is modelled by deactivating a part of the motor cortex in the model. Since stroke is introduced at the level of motor cortex which is the hub of integration of the inputs from all the other components (observed in eqn **18**), presence of stroke at this level will have a direct influence on the performance of the corresponding arm, and to some extent even in the other arm due the coupling between the two motor cortices in the model.

### Unimanual reaching

Several control and stroke studies are performed using the model to compare and quantify the effects of stroke. The effect of stroke in the model on the reaching behaviour of the arm is evaluated in terms of two performance indices: PRV and reaching error. The workspace of the arm that comprises of all the reachable target locations is mapped under both stroke, as well as post intervention paradigms. Under stroke conditions, there is an evident decrease in the number of targets the arm can reach in its workspace because of the deactivation of certain nodes in the motor cortex due to which it fails to form representations for those goal positions (Fig. 5B). Post intervention, by retraining the connections between the PC and MC that corresponds to the representations lost due to lesion and MC and MN that accounts for the specific configurations of the arm that were absent under stroke conditions, the ability of the arm to reach the targets in its workspace that were previously unreachable, improved. Thus the current model accounts for the recovery of the arm after intervention and also demonstrates the potential to explain functional plasticity in the brain (Cheatwood, Emerick et al. 2008).

The arm’s movement when it reaches the target is analyzed and the trajectories are recorded. During the initial stages of reaching the target (between trials 1 and 20) where the arm accomplishes the target, the trajectory is observed to have high path variability (Muralidharan, Mandali et al. 2018). Early stage of training in the model is characterized by a low contribution from pre-frontal cortex and high basal ganglia contribution. The exploratory drive originating from the complex and chaotic dynamics of the STN-GPe system of the BG seems to account for high path variability in this stage. The trajectories in the later stages of training (between trials 41 and 50), become smoother with reduced path variability (Fig.3D). In late stage training, the contribution from the BG is reduced, with lesser influence from the STN-GPe system, and thereby lesser exploration. These features perhaps explain the reduced path variability in later stages.

The velocity of the arm as it reaches the target, shows the characteristic bell-shaped curve profile, though the PRV of the arm under stroke conditions is observed to be lesser than the PRV under normal conditions (Fig. 6C). Thus, the introduction of stroke in the model introduces a temporal delay and reduces the performance of the arm in reaching tasks as is observed in the classic case of upper extremity hemiparesis (Rose and Winstein 2005). These studies effectively indicate the model’s potency to simulate unimanual reaching behaviours under normal and stroke conditions.

### Bimanual Reaching

The paper focuses on modelling and simulating the task setup of Rose et al (2004) for bimanual training as a prospective rehabilitation strategy for stroke.

The primary performance index analyzed in this study is the velocity profile of both the arms when they carry out reaching tasks. It is found that during the symmetric aiming task (in which the distance of the targets from its respective arms are the same) under unimanual conditions, where both the arms are independently allowed to accomplish their tasks, the PRV of the non-paretic arm (left arm) is higher than the PRV of the paretic arm (right arm). Conversely, when the arms performed the same reaching tasks under bimanual conditions, the paretic arm showed an increase in its PRV and the non-paretic arm showed a significant decrease (Fig. 8). This result suggests that the non-paretic arm aids the paretic arm in moving towards its target, when the arms perform the task under bimanual conditions.

A second task is performed by introducing an asymmetry in the target locations. Here again there are two cases. In the first case, in the so-called congruent aiming task, the paretic arm has a near target and the non-paretic arm has a far target. It is observed that when the arms performed the task under both unimanual and bimanual conditions, there seemed to be no significant change in their PRVs: the PRV of the non-paretic arm is found to be higher than the paretic arm in both the cases (Fig. 9). It may be inferred that a relatively easier performance target for the paretic arm has an insignificant influence on its PRV and hence does not contribute to any effective improvement of the arm.

The second case is known as the incongruent aiming task. Here, the paretic arm has a far target whereas the non-paretic arm has a near target. The PRV’s were recorded in the same fashion as before. It is observed that under unimanual conditions, the non-paretic arm has a higher PRV than the paretic limbs as is the normal case. But a surprising change is observed under bimanual conditions, where there is a significant increase in the PRV of the paretic arm accompanied by a significant decrease in the PRV of the non-paretic arm (Fig. 10). It can be deduced from these changes in the PRV, that using a more challenging target for the paretic arm helps the paretic arm’s functional recovery (under bimanual conditions) more than using an easy one. The model is able to capture all of the abovementioned performance variations mentioned in the empirical study (Rose and Winstein 2004).

### Selecting Rehabilitation Strategies to treat Stroke

Stroke rehabilitation requires a well-rounded understanding of the post-stroke effects or loss of functionality that patients suffer. Although interventions begin within the first 48 hours of stroke manifestation, only 60% of people with hemiparesis have received functional independence in activities of daily living (ADL) (Nakayama 1995; Patel, Duncan et al. 2000). Hence, a comprehensive and customized treatment strategy that would vary from patient to patient is required for effective treatment. Progressively training components of goal-oriented tasks by reinforcing behaviour using specified learning networks or in other words, physical training has proved to be a go-to and an effective strategy implemented in stroke rehabilitation (Dobkin 2004) followed by other techniques that have emerged such as robotic arm training (Fasoli, Krebs et al. 2004), virtual reality approaches (Jang, You et al. 2005; Chang, Tung et al. 2007; Turolla, Dam et al. 2013) etc. The motivation behind our paper is to study these conventional training therapies using a computational model and allow the model to assess the relative merits of different therapies. To begin with, we have focused on three different rehabilitation strategies that fall under physical training therapies: Unimanual reaching tasks (URT), bimanual reaching tasks (BRT) and Constraint Induced Movement Therapy (CIMT).

CIMT is modelled by arresting the non-paretic arm (left arm in this case) in a specific configuration (this is similar to constraining the arm using a sling as observed in the experiments). The right arm (paretic arm) thus receives a constant motor cortex activity from the constrained arm modulated by the coupling strength (ε_R_ or ε_L_). Different comparison studies are performed between these rehabilitation strategies. The first study was performed by introducing *acute* stroke in the model and measuring the reaching error of the arm. Reaching error is defined as the closest distance to which the arm gets to the target location when it performs the task. This analysis is carried over different lesion sizes to assess the efficacy of the chosen rehabilitation strategy as a function of the size of lesion in the motor cortex. It is found (Fig. 11A) that until lesion size of 3×3 (which denotes small lesion sizes) all three rehabilitation strategies (unimanual, bimanual and CIMT) prove to be effective, and their corresponding reaching errors are zero or approximately zero‥ Beginning from a lesion size of 4×4 until 6×6, it can be seen that the value of reaching error is zero for bimanual but has a significant value under unimanual and CIMT conditions, thus making bimanual the most sought after strategy in this lesion bracket. Interestingly, the opposite is observed for a lesion size of 7×7 with bimanual reaching error being higher than the unimanual and CIMT value. However the difference in the value is not that significant.

The second study is carried out under *chronic* stroke conditions. For 1×1 lesion, both CIMT and bimanual training prove to be effective strategies. But from lesions of size 2×2 to 5×5, higher reaching errors in both unimanual and CIMT trainings make it a non-preferred strategy for relatively smaller lesion sizes. On the other hand, bimanual training shows a reaching error of zero or approximately zero, until a 5×5 lesion size but spikes suddenly at 6×6 thereby making it an unreliable intervention strategy at greater lesion sizes (Fig. 11B). Thus we conclude that unimanual reaching and/or CIMT are more preferred rehabilitation strategies for greater lesion sizes in the motor cortex.

## Conclusion

Although the model gives rise to some useful predictions with respect to developing customized therapies for stroke, insufficient data\knowledge about the cortical and sub-cortical connectivities still remain a challenge to simulate biologically plausible coupling within different regions of the motor network. However, the model intelligibly accounts for the inter-hemispherical connections by incorporating the coupling factor (ε) at the level of motor cortices. This is justified because the motor cortex component receives input from all the other components in the model and integrates it, dynamically influencing the performance of the system.

The model also shows the potential to be developed into a more comprehensive system for targeted reaching by appending the existing model with unified reward and value function. 3-D reaching tasks can also be performed and more realistic motor and somatosensory maps can be generated using LISSOM (Laterally Interconnected Synergistically Self Organising Maps). We also aim to develop a more comprehensive framework of the motor system by augmenting the existing model with a cerebellum component. Thus the current model holds immense potential to be developed and used as one of the key clinical tools employed for stroke rehabilitation.

## Methods

To examine the after effect of stroke on upper extremity function, we use a cortico-basal ganglia model (Muralidharan, Mandali et al. 2018) capable of performing simple bimanual reaching movements. In essence, the model has two semi–independent systems, each capable of driving reaching movements in a single hand. Each system has an outer loop corresponding to the cortical loop and an inner loop corresponding to the basal ganglia loop. The outer loop consists representations of two cortical modules – the *proprioceptive cortex* and the *motor cortex*, - and another module representing the spinal cord. The combined cortico-basal-ganglia system controls a simple two-link arm model. A pair of systems, corresponding to the left and right hemispherical systems, is coupled at the level of the respective motor cortices, controlling their respective arms. There exists a communication between the two motor cortices by means of a coupling factor (ϵ), reflecting inter-hemispherical connectivity. The proposed architecture is described in greater detail as follows:

### A. Model architecture

The model architecture comprises of an outer sensory motor-cortical loop and an inner cortico-basal ganglia loop. The outer loop consists of the two-link kinematic arm, proprioceptive cortex (PC), motor cortex (MC) and motor neurons (MN) that innervate the two muscle pairs of the two link arm. The inner loop comprises of the basal ganglia (BG) and its components such as the striatum, globus pallidus external and internal segments (GPe and GPi), the subthalamic nucleus (STN) and the thalamus (Fig.12).

**Fig.12:**
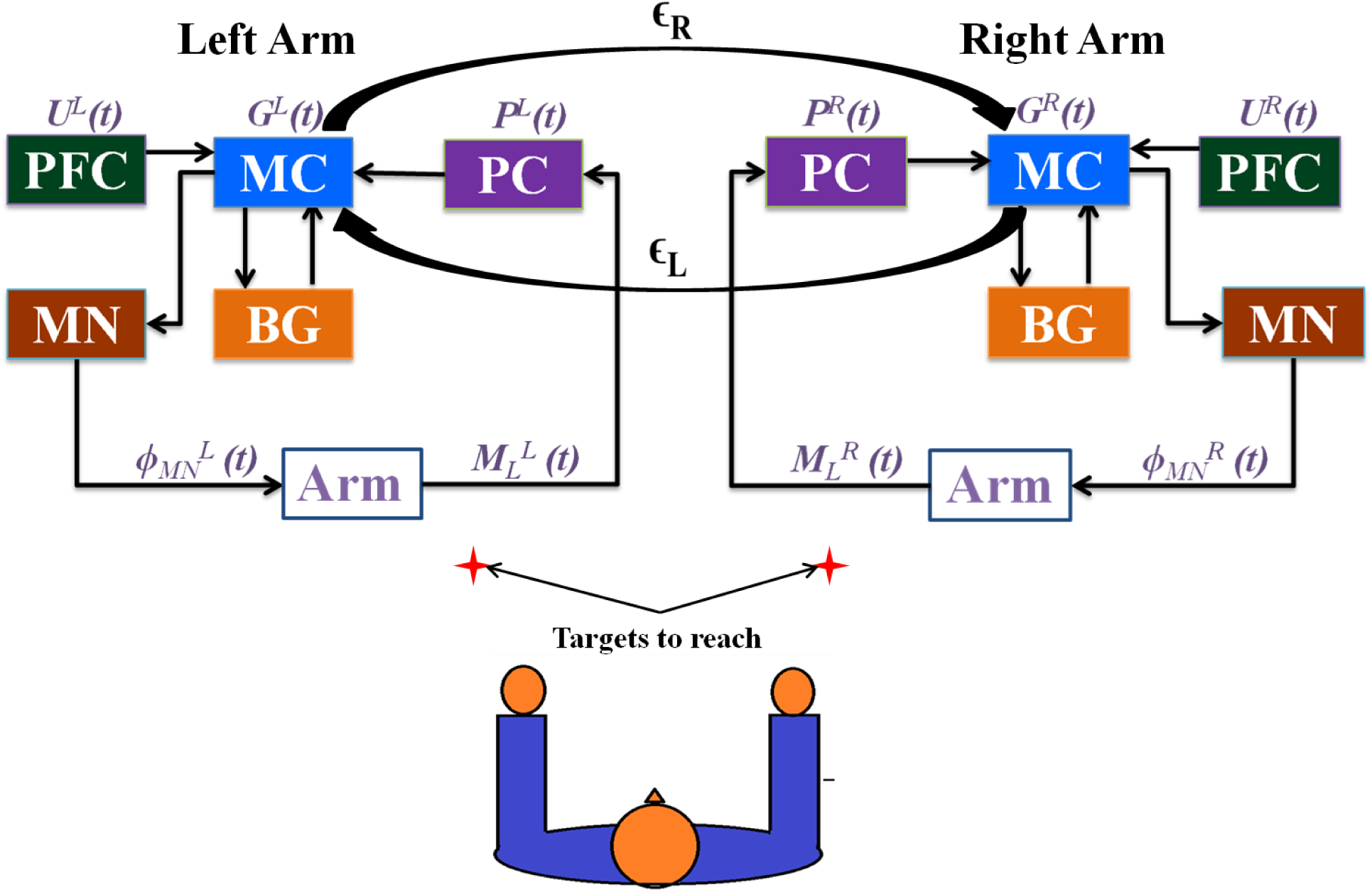
The cortico-basal ganglia model for bimanual reaching. The architecture is designed to have two loops, a sensory-motor “outer” loop and the cortico-basal ganglia “inner” loop. The motor cortex receives projections from higher frontal areas which in the model is the Prefrontal cortex. The size of the neuronal sheet used for each module in the model is 15×15.

A detailed description about every component with respect to single arm module is as follows:

#### 1. The Sensory motor cortical loop

##### 1.1 Arm model

The two link kinematic arm is composed of two joints where each joint is controlled by an agonist (Ag) and an antagonist (An) muscle pair. Each pair is in turn innervated by a pair of motor neurons represented in the form a four dimensional vector ϕ^MN^(t). The activations of the innervated muscle pairs are then used to obtain the shoulder and the elbow joint angles 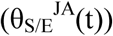 given by the equations:

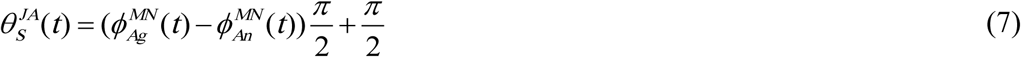

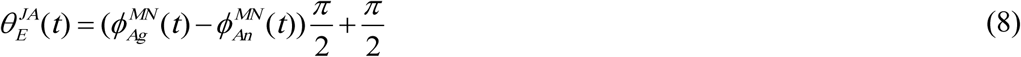

The measures of the joint angles define the range of movements of the arm over the 2D workspace which contains a given set of targets. Following this, the lengths (*μ*^E^ and *μ*^S^) of each muscle is calculated from the joint angles by using the following equations:

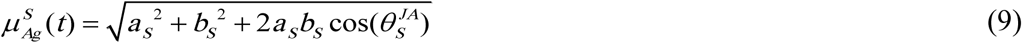

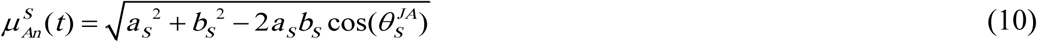

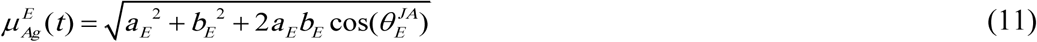

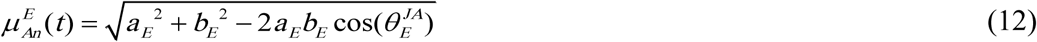

A sensory (proprioceptive) map of the arm is subsequently generated from the four dimensional muscle length vector 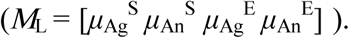 The end effector location of the arm 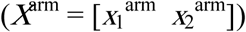 is computed from:

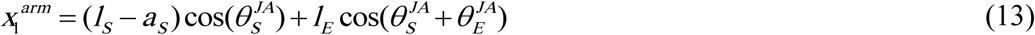

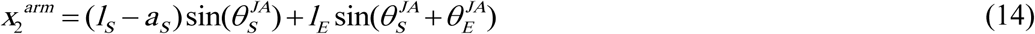

##### 1.2 Formation of the sensory map

The sensory map of the arm is generated by the PC, which is modeled as a Self-Organizing Map (SOM) (Kohonen 1990) of size *N*_PC_ x *N*_PC_. The muscle length vector (*M*_L_(t)) is used as the feature vector to train the PC SOM. The activation of a single node *i* in the PC is given by:

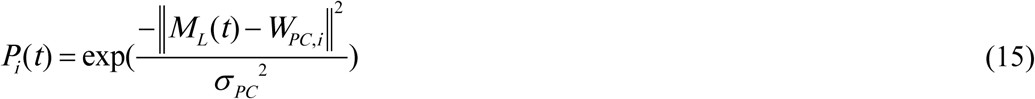

##### 1.3 Formation of the motor map

The motor cortex comprises of a 2D sheet of neurons of size *N*_MC_ x *N*_MC_ and is modeled as an amalgamation of SOM and Continuous Attractor Neural Network (CANN) (Trappenberg 2005) to account for the characteristic of low dimensional input data representation and dynamics exhibited in such cortical areas. A dynamic model like CANN is employed to facilitate the integration of multiple afferent inputs received from the PC, the BG and the Pre-Frontal Cortex (PFC).

The lateral connectivity in the CANN model is characterized by short range excitation and long range inhibition whose dynamics are defined by the weight kernel 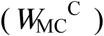 given by,

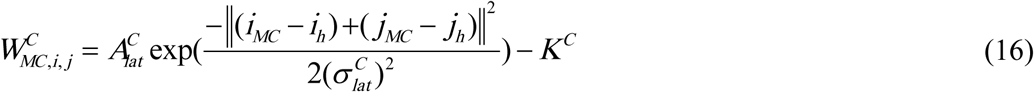

where,

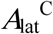 is the strength of the excitatory connections

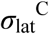 is the radius of the excitatory connections and

*K*^C^ is the global inhibition constant

[*i*_MC_, *j*_MC_] are the locations of the nodes in MC, [*i*_h_, *j*_h_] corresponds to the central node.

The output from PC, a matrix of size *N*_PC_ x *N*_PC_, is converted into a vector of size 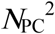 x1 and given as input to the SOM part of MC for the development of motor map of the arm. There exists all-to-all connections from the muscles of the arm to PC and PC to MC. The MC SOM is trained using the standard SOM algorithm (Kohonen 1990). The activation of a node *i* in the SOM part of the MC is given by:

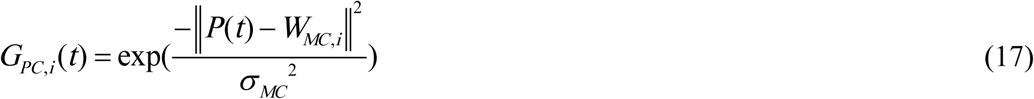

where,

*W*_MC,i_ is the weight connection between the PC and the *i*^th^ node of the SOM part of MC *σ*_MC_ is the width of the Gaussian response.

The outputs from PC (G_PC_), BG (G_PC_) and PFC (G_PFC_) are then presented as input to the MC CANN. The input equation is given by:

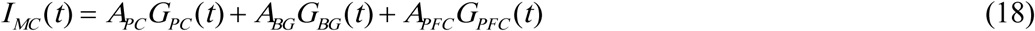

where, *A*_PC_, *A*_BG_, *A*_PFC_ are the respective gains of the PC, BG and PFC networks.

With these inputs, the activation dynamics of the MC is given by:

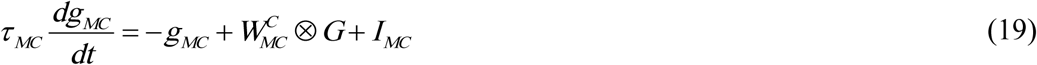

where *g*_MC_ is the internal state of the MC neurons.

The output MC activity (*G*(*t*)) is given by,

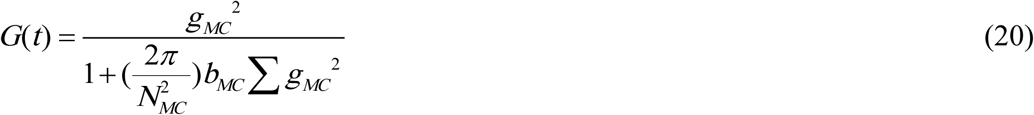

The MC neurons project into the four motor neurons whose activation is then given by,

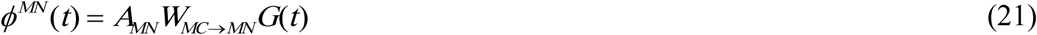

To close the sensory motor loop, the MC to MN connections are trained in a supervised manner. To begin with, we give direct external input to MN, which may be considered as the desired MN activation 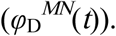 This input will in turn produce a movement in the arm which in turn produces responses in the PC and MC in that sequence. The new MC activation presents a new input to the MN module. Now the desired MN activation 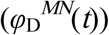 and the actual MN activation (*φ^MN^*(*t*))) obtained after traversing over the cortical loop must be ideally the same. But in an untrained cortical loop there will be a difference. The difference between the desired and actual MN activations is used to train the connections between the MC and the MN (*W*_MC→MN_) layer:

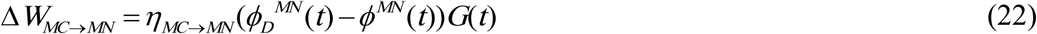

where, Δ*W*_*MC*→*MN*_ corresponds to weight updation and *η*_*MC*→*MN*_ is the learning rate.

Since the model is bimanual, the motor cortex of one arm is connected to the motor cortex of the other by means of a coupling factor (ϵ). Thus, a connection is established between the two hemispheres at the motor cortical level. To study upper limb hemi paretic stroke paradigms, we introduced a “lesion” in the right motor cortex by nullifying the activity of a part of the MC.

#### 2. The Basal Ganglia

The fundamental principle of BG operation is reinforcement learning where the BG learns to choose optimal actions based on a reward feedback mechanism (Chakravarthy and Balasubramani 2015). Hence the BG drives the arm via the motor cortex and enables the arm to reach the target. The process of action selection is guided by the presence of a value function within the striatal module of the BG. Thus, the arm will learn to choose the action which brings the greatest increase in value. In the model, the value function codes for the error between the desired goal position (*X*^targ^) and actual end effector position (*X*^arm^) in terms of distance. The BG output then performs a stochastic hill climbing over the value function to search for the maximal value. The value is calculated by,

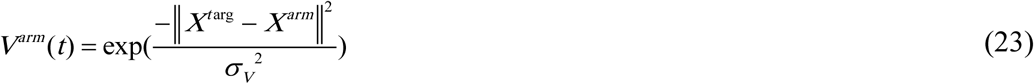

where,
***σ***_V_ defines the spatial range over which the value function is sensitive for that particular target.

We then determine the value difference signal (*δ*_v_) which regulates the switching between direct and indirect pathways.

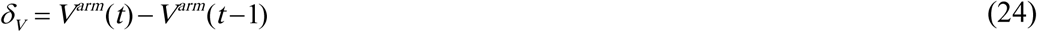

The switching happens due to the modulation of the responses from the striatal Medium Spiny Neurons (MSN). This is represented as:

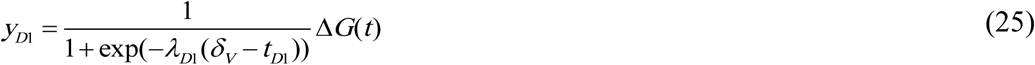

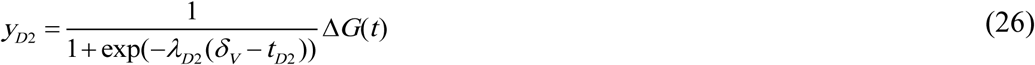

where, Δ*G*(*t*) refers to the difference vector represented as difference in motor cortical activity; *y*_D1_ and *y*_D2_ represent the outputs of D1R- and D2R-expressing Medium Spiny Neurons (MSNs) of the striatum. *λ*_D1_ and *t*_D1_, *λ*_D2_ and *t*_D2_ are the gains and the thresholds of the direct and indirect pathway respectively.

The dynamics of the STN-GPe system influenced by *y*_D2_ is given by,

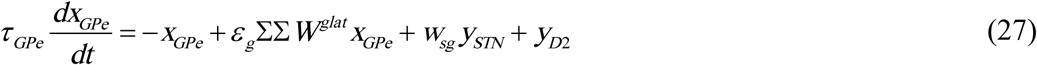

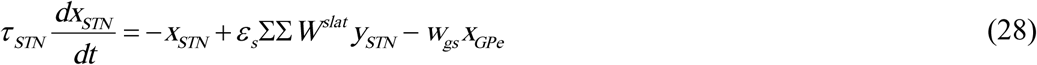

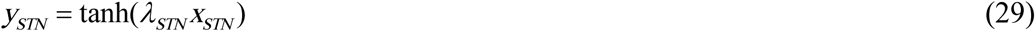

where,

*W^slat^* and *W^glat^* are lateral weight connections with connection strengths *ϵ*_s_ and *ϵ*_g_ within STN and GPe respectively

*w_sg_* and *w_gs_* are the weight parameters that control the connection strengths.

***τ***_STN_ and ***τ***_GPe_ are the respective time scales of STN and GPe.

*λ*_STN_ controls the STN output by controlling the slope of the sigmoid.

***τ***_STN_ and ***τ***_GPe_ are the time scales of STN and GPe respectively.

The Gaussian neighborhood of the lateral weight connections is given as

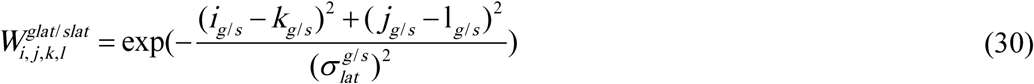

where, 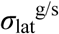 is the spread of the lateral connections respectively for the STN-GPe network.

The exploratory behavior of the arm is attributed to the uncorrelated oscillations of the STN layer which are produced as a consequence of low striatal inputs. This is due to the formation of excitatory-inhibitory neuron pools by the STN-GPe system constituting the indirect pathway. Such excitatory-inhibitory pairs of neuronal pools are known to exhibit complex oscillations (Chakravarthy and Balasubramani 2015).

The output signal of the direct pathway from the D1R-expressing MSNs in the striatum is integrated with the STN output in the GPi as follows:

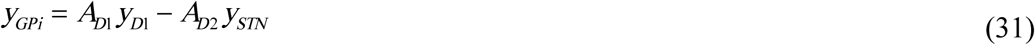

This output from the GPi is then presented to the thalamus, also modeled as a continuous attractor neural network (CANN).

#### 3. Representation of goal location

Goal of a movement is considered to be represented in the PFC, thereby making it the source of a motor command (Matsumoto, Suzuki et al. 2003; Asplund, Todd et al. 2010). Hence, in our model the PFC is used to convey information regarding the goal position to the motor cortex. The PFC is modeled and trained as a SOM, with weights *W*_PFC_. The locations accessible by the arm in its 2D workspace are given as input feature vectors to train the SOM. The activation in the PFC corresponding to a particular target location is given by:

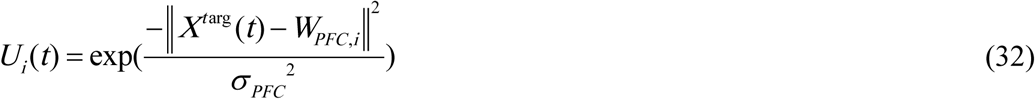

The PFC to MC weight connections (*W*_PFC→MC_) are trained whenever the arm reaches the target. If *G*_PFC_ is the activity that is induced in the MC due to PFC activation and *G*_targ_ is the MC activity that facilitates the arm to reach its target, then the training of weights between PFC and MC is given as:

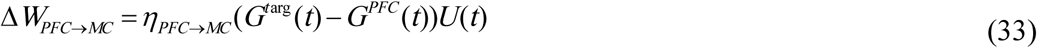

At the outset, learning occurs as the result of slow movements of the arm governed by the BG. However, in later stages, learning occurs due to fast movements dominated by the cortical loop. Hence PFC contribution increases as a function of number of trials.

### B. Simulation of aiming conditions

In the model, each arm had its respective target placed at locations in a manner identical to that of the experiment conducted by Dorien and Rose (Rose and Winstein 2004). In the experiment, the subjects were asked to aim and hit the switches (targets) using either one arm or both arms in response to an LED signal. Based on target locations, the tasks given to the subject/model to perform were classified into:

i. Symmetric/Equidistant aiming
ii. Congruent aiming
iii. Incongruent aiming

All three tasks are implemented in the model, in both unimanual and bimanual conditions. In unimanual condition, the arms act independently (ϵ = 0) and in the bimanual reaching condition, they reach their respective targets simultaneously (ϵ < 0 and ϵ > 0) (Table 1). Congruent aiming task involves movement of the paretic arm to a nearby target (“near” target) with the normal arm aiming at a far off target (“far” target). The distance between the near target and the subject was half the distance between the far target and the subject. Similarly in incongruent aiming, the paretic arm had to reach the far target while the non-paretic arm moves toward a near target. These tasks are used to study the effect of reaching distance and aiming condition on the performance of the paretic arm.

**Table 1.** Parameters used in the model.

| Arm Kinematics |  | Learning Rates |  | Coupling Factor |  |  |  |
| --- | --- | --- | --- | --- | --- | --- | --- |
| $a_S$ | 0.04 | **MC→MN Net** | | Symmetric Aiming | | | |
| $b_S$ | 0.07 | $\eta_{MC \rightarrow MN}$ | 0.1 | $\varepsilon_r$ | -0.5 | $\varepsilon_l$ | 0.5 |
| $a_E$ | 0.03 | **PFC→MC Net** | | Congruent Aiming | | | |
| $b_E$ | 0.08 | $\eta_{PFC \rightarrow MC}$ | 0.1 | $\varepsilon_r$ | -0.2 | $\varepsilon_l$ | 0.89 |
| $l_S$ | 0.3 | | | Incongruent Aiming | | | |
| $l_E$ | 0.3 | | | $\varepsilon_r$ | -0.2 | $\varepsilon_l$ | 0.8 |

Besides the implementation of unimanual and bimanual conditions, we have also implemented in the model, Constraint induced movement therapy (CIMT), an intervention that has shown to be effective in restoring functionality of the stroke affected limb (Taub, Lum et al. 2005; Taub, Uswatte et al. 2006). The objective of this part of the modelling study is to investigate the effect of CIMT on stroke rehabilitation and understand its limitations. CIMT involves forced usage of the affected arm to perform tasks while actively restraining the unaffected arm by means of a sling or a splint. In the model, this is achieved by maintaining the unaffected arm (left) in a fixed initial configuration and allowing the paretic arm (right) to perform the task. Fixing the arm in a particular configuration ensures the imposition of a “constraint-induced” framework.

### C. Lesion study

To analyse the impact of the size of lesion in the MC on the performance of the paretic arm under different aiming conditions, lesions of varying size ranging from 1×1 to 7×7 are introduced in the MC. The coordinates of the center of the lesion (i_les_, j_les_) is selected such that activation of the MC near (i_les_, j_les_) places the arm at the goal location. The center of the lesion is fixed while the size is varied. This study is conducted for three aiming conditions namely unimanual, bimanual and CIMT. The arm is trained under all three conditions and is tested using symmetric targets. The lesion is centred on the (i, j) ^th^ node and is expanded from a size of 1×1 to 7×7. The reaching error of the arm is computed for every lesion size as function of the distance between the actual target and end effector positions at the end of every trial.

## Contribution Of Authors

All authors contributed equally to the work. Rukhmani Narayanamurthy, Samyukta Jayakumar and Sundari Elango performed coding, analysis of the model and manuscript preparation. Vignesh Muralidharan performed designing the model, coding, analysis of the model and manuscript preparation. V. Srinivasa Chakravarthy performed designing the model and manuscript preparation.

